# An Exploratory EEG Comparison of Good and Bad Readers

**DOI:** 10.1101/2024.12.17.628834

**Authors:** María José Álvarez-Alonso, Cristina de la Peña, Ricardo Scott

## Abstract

Reading is a crucial skill for critical thinking and cognitive development, yet many individuals experience difficulties that impact their daily lives. This study examines the neurophysiological differences between proficient and struggling readers using electroencephalography (EEG). The analysis focuses on the alpha (8-12 Hz) and beta (13-30 Hz) frequency bands, which are prominently involved in cognitive and language-related processes, although other frequency bands also contribute significantly to language comprehension and cognitive control. Eighteen participants (9 good readers and 9 poor readers) were selected based on standardized reading assessments. Resting-state EEG recordings were collected with a 64-channel system while participants were at rest with their eyes closed. After preprocessing to remove artifacts, the power spectra were analyzed, emphasizing relative power and alpha peak activity. Functional connectivity was measured using the corrected imaginary part of the phase-locking value (ciPLV), ensuring accuracy by minimizing volume conduction effects. Results revealed that good readers displayed increased beta power in frontal regions and enhanced synchronization within fronto-central-parietal networks compared to poor readers. Alpha band activity showed complex associations with factors such as age, reading skills, and verbal fluency, indicating nuanced relationships between neural development and literacy. The heightened beta activity in good readers is consistent with its role in cognitive control and language processing, while their stronger network connectivity suggests more efficient neural communication. These findings provide valuable insights into the neural basis of reading proficiency and emphasize the importance of distributed brain networks in skilled reading. Future research should replicate these results with larger samples and longitudinal designs to better understand the neural mechanisms underlying literacy development.

## INTRODUCTION

In contemporary society, characterized by rapid change and complexity, the ability to read proficiently is essential for effective decision-making and critical thinking. Reading enables individuals to interpret and synthesize complex information, serving as a foundational skill for higher-order cognitive processes. van Gelder (2005) highlights that critical thinking builds upon such foundational skills, as it requires the ability to evaluate and integrate diverse sources of information. However, Willingham (2007) argues that critical thinking is not merely a deployable skill but a form of reasoning that can be inconsistently applied, even by experts. This suggests that while reading comprehension is a necessary foundation, developing robust critical thinking abilities involves additional factors, underscoring their intricate relationship.

Many individuals, despite completing formal education, struggle with adequate reading skills, impacting their cognitive development and daily functioning. Reading difficulties can manifest as challenges in decoding words, fluency, and comprehension. The significance of literacy extends beyond basic decoding abilities; it encompasses higher-order skills essential for navigating complex information environments. In fact, UNESCO (2005) emphasizes that literacy includes the capacity to synthesize information and think critically. This broader definition aligns with current research on the multifaceted nature of reading comprehension and its role in cognitive development.

Importantly, Morais and Kolinsky (2020) highlight that these difficulties are not indicative of a lack of intelligence but rather stem from specific neurocognitive differences. This perspective shifts the focus from general cognitive ability to targeted neurocognitive processes involved in reading. Research into media literacy underscores its potential to foster critical engagement with information, particularly in combating misinformation and fostering analytical thinking. Bulger and Davison (2018) emphasize that media literacy, as a skillset rooted in critical inquiry, empowers individuals to evaluate, interpret, and respond to complex media messages. This approach aligns with broader cognitive competencies, such as decoding, fluency, and comprehension, highlighting that challenges in these areas are often less about cognitive deficits and more about the need for targeted educational interventions.

In light of these considerations, this exploratory study aims to explore the neurophysiological differences between proficient and deficient readers using electroencephalography (EEG). By examining power spectrum analysis and functional connectivity patterns, we seek to elucidate the neural mechanisms underlying reading proficiency, building on the growing body of neuroscientific research in this field (Price y Devlin, 2011; Bedo et al., 2021).

## METHODS

### Participants

Eighteen participants were recruited for this study and divided into two groups based on their reading proficiency: high-readers (n=9) and low-readers (n=9). The classification was conducted using the PROLEC-SE-R Reading Process Assessment Battery (Cuetos et al., 2016), a standardized tool designed to evaluate lexical, syntactic, and semantic processes in reading. This approach followed established protocols from previous studies (Álvarez-Alonso et al., 2021; Horowitz-Kraus et al., 2014). In addition to reading proficiency, participants were assessed on cognitive tasks using subscales of the Wechsler Adult Intelligence Scale - Fourth Edition (WAIS-IV) (Wechsler, 2008), specifically targeting attention, coding, verbal reasoning, and fluency. To further analyze attentional processes, the d2 Test of Attention (Brickenkamp & Zillmer, 1981) was administered. Demographic factors and school grading levels were also collected to contextualize the findings.

Neurophysiological data were acquired using a portable 64-channel EEG amplifier with gel-based electrodes from ANT NEURO - eego™ my lab (ANT NEURO, Germany) at a sampling frequency of 500 Hz. The electrodes were placed according to the extended 10-20 system positions (e.g., Fp1, Fpz, Fp2) using high-quality Ag/AgCl sensors. The reference electrode was positioned at Cpz, and offline re-referencing was conducted to average reference. Sensor impedances were maintained below 10 kΩ. Each participant underwent a 5-minute resting-state EEG recording with eyes closed while seated comfortably, following established protocols for resting-state EEG studies (Klimesch, 2012).

### Data Preprocessing

EEG data were examined for ocular, muscular, and jump artifacts using the OHBA Software Library (OSL). Artifact-free data were segmented into four-second epochs and band-pass filtered between 2-45 Hz with 2-second padding, adhering to best practices in EEG data preprocessing (Bedo et al., 2021).

### Power Spectrum and Functional Connectivity Analysis

The power spectrum for each EEG channel was computed using Fast Fourier Transform with Hanning windows smoothed at 0.25 Hz. Relative power was calculated by normalizing total power in the 2-45 Hz range. Specific alpha band power (8-12 Hz) was also analyzed, given its established role in cognitive processing (Klimesch, 2012). Functional connectivity was assessed using the corrected imaginary part of the phase locking value (ciPLV), which evaluates phase synchronization across sensor pairs (Bruña et al., 2018; Lachaux et al., 1999). This method was chosen for its robustness against volume conduction effects. Statistical analyses were performed using Mann-Whitney tests for independent samples to compare group differences. Graph measures such as topology, strength, efficiency, modularity, and centrality were computed using the Brain Connectivity Toolbox (BTC), following established network neuroscience approaches (Rubinov & Sporns, 2010).

## RESULTS

### Power Spectrum

Figure 1 shows the power spectrum for both groups divided by reading proficiency. The overall power spectrum (left panel) demonstrates a similar pattern for both groups, with no statistically significant differences observed. However, a slight reduction in alpha peak power (8–12 Hz) can be visually identified in the high-proficiency readers compared to the low-proficiency readers. This pattern is consistent in the occipital sensors (right panel), where the high-proficiency group again displays a marginally lower alpha peak. These observations are based on descriptive analysis and lack statistical significance, as determined by the applied tests.

**Figure 1.**
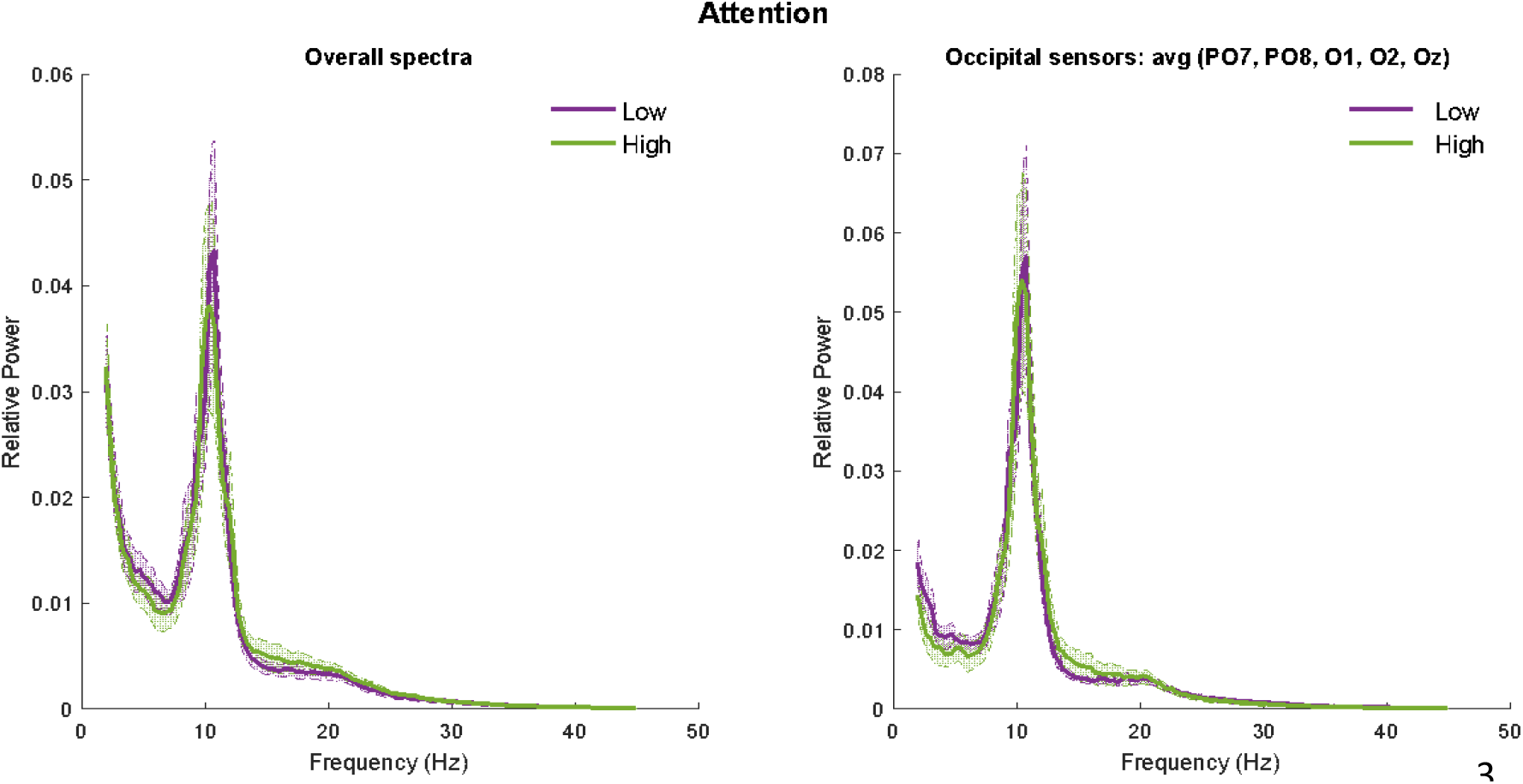
Left-panel: Overall power spectrum (mean of all channels) for the high attention group (green) and low attention group (purple). Right-panel: Occipital power spectrum (mean of occipital sensors) for the high attention group (green) and low attention group (purple).

Figure 2 shows the distribution of beta power across the scalp for both groups, as illustrated in the topographical maps (upper panel). These maps highlight a consistent pattern of beta activity between groups, with noticeable regional variations. In the lower panel, the bar graphs display statistically significant differences (p < 0.05) in beta power between the two groups, specifically in sensors AFz and F3. These results suggest localized differences in beta activity, which may reflect variations in cognitive processing related to reading proficiency. Low reading and high reading are referred to low and high forms of reading in the test.

**Figure 2.**
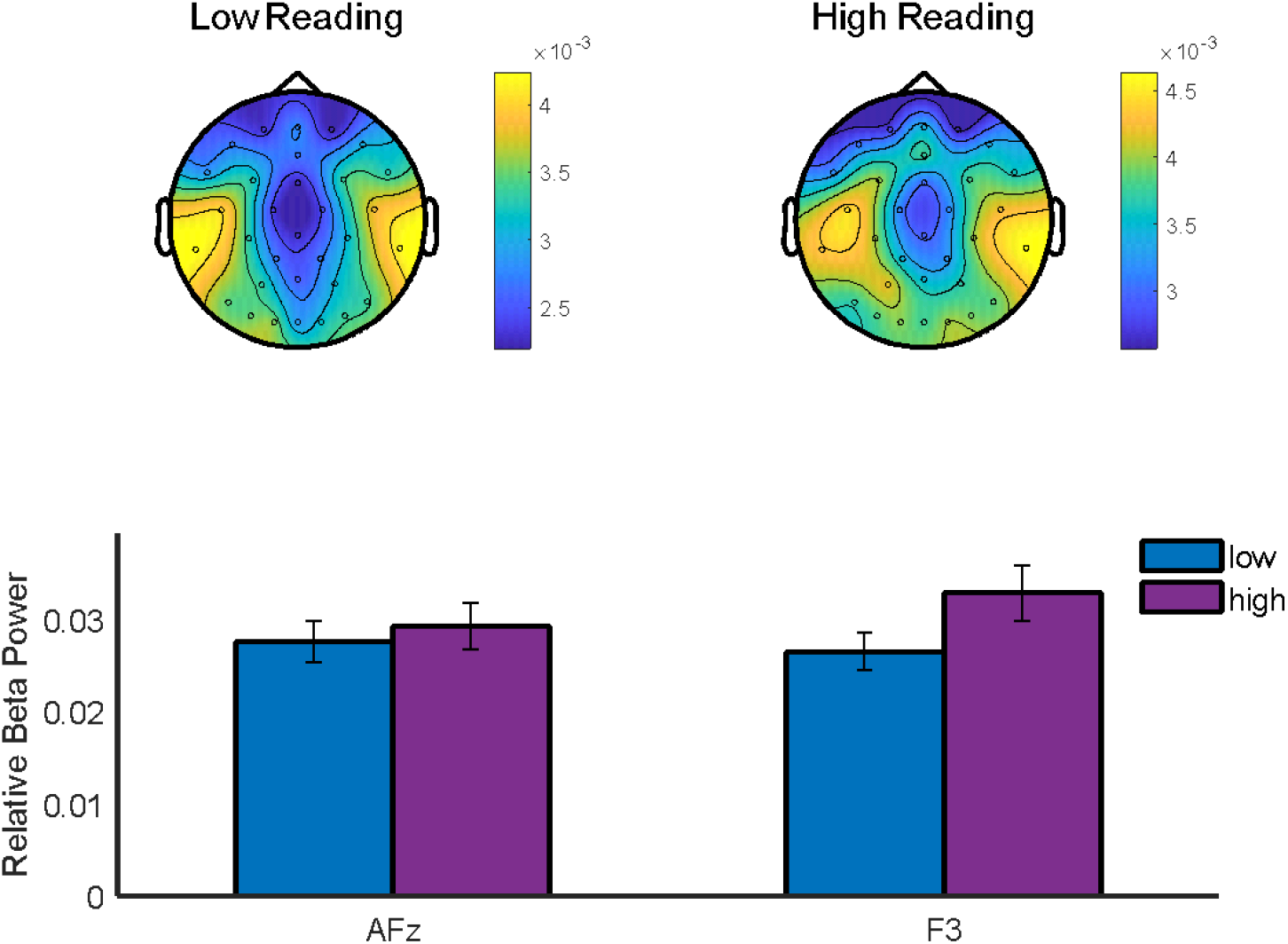
Beta Band Power. Upper-panel shows alpha band distribution for the two groups in the attention condition. Lower-panel shows the statistically significant differences (p<0.05) in sensors AFz and F3 for both groups.

### Functional connectivity

Figure 3 illustrates the differences in functional connectivity between the high and low reading proficiency groups. The upper panels display statistically significant links between electrode pairs when comparing the two groups. Notably, the high-proficiency group exhibits increased connectivity compared to the low-proficiency group (p < 0.01). The bar plots in the lower panels quantify these differences, showing higher average beta-band functional connectivity across multiple electrode links in the high-proficiency group. These results suggest enhanced neural coordination in the high-proficiency group, which may reflect more efficient information integration during cognitive tasks.

**Fig 3.**
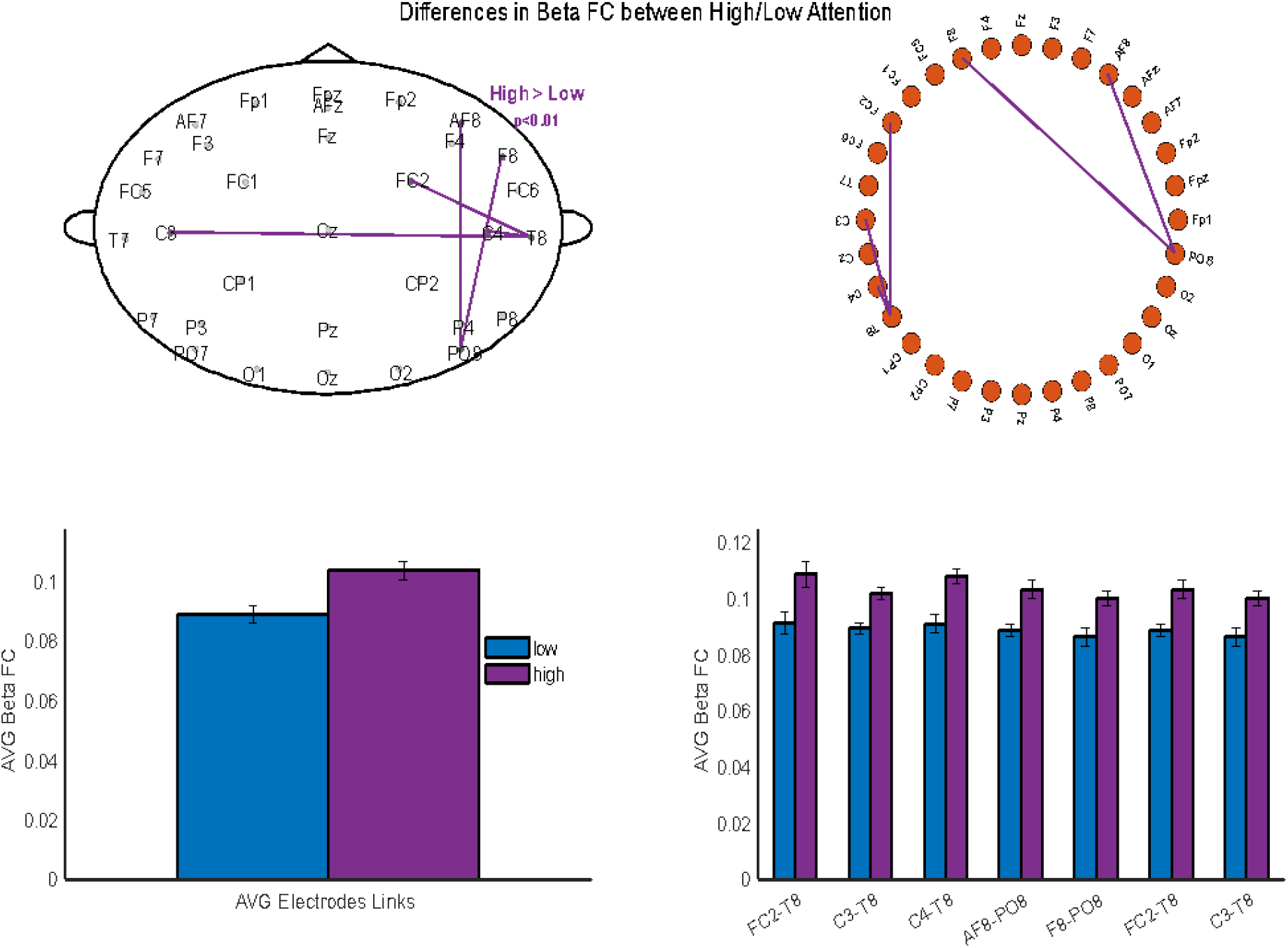
Functional Connectivity Analysis. Upper-panel shows the connectivity distribution (high>low – p<0.01) over the scalp. Lower-panel shows the mean of all connectivity links and the mean FC for each link found.

### Correlation analyses

This study investigated the neurophysiological correlates of reading ability by comparing high and low reading groups using electroencephalographic (EEG) measures and their associations with various cognitive and demographic factors. The results revealed distinct patterns between the two groups, particularly in the alpha and beta frequency bands. In the low proficiency reading group, significant correlations were observed between age and beta power in the F3 electrode (r=0.783, p=0.021), school reports grading and alpha peak (r=0.731, p=0.040), and phonological verbal fluency (A) with both beta participation coefficient (r=0.761, p=0.028) and beta power in the F3 electrode (r=-0.798, p=0.018). Additionally, results in the coding task showed negative correlations with beta participation coefficient (r=-0.880, p=0.004) and alpha band power (r=-0.807, p=0.015), while CD2 was negatively associated with alpha betweenness (r=-0.778, p=0.014) and beta power in the O1 electrode (r=-0.715, p=0.030). The high reading group exhibited more complex patterns of correlations, particularly in the alpha band. Age demonstrated both positive and negative correlations with alpha band measures, including betweenness (r=0.669, p=0.034), participation coefficient (r=-0.814, p=0.004), and eigencentrality (r=-0.674, p=0.033). Reading ability was positively correlated with alpha betweenness (r=0.640, p=0.046) and negatively with alpha participation coefficient (r=-0.791, p=0.006). Phonological verbal fluency measures (F, A, and S) consistently showed positive correlations with alpha participation coefficient and eigencentrality, while negatively correlating with alpha strength functional connectivity (FC), betweenness, clustering, and global efficiency. Similarly, TAD2, OD2, and COND2 displayed positive correlations with alpha participation coefficient and eigencentrality, and negative correlations with alpha betweenness. These findings suggest distinct neurophysiological profiles for high and low reading groups, with the high reading group demonstrating more complex network characteristics, especially in the alpha band. The observed associations between cognitive measures and various network metrics in the high reading group highlight the intricate nature of neural processes underlying advanced reading abilities. In contrast, the low reading group showed fewer but still significant correlations, primarily in the beta band. These results contribute to our understanding of the neural underpinnings of reading ability and emphasize the importance of considering multiple frequency bands and network measures when studying cognitive processes related to reading proficiency.

## DISCUSSION

The analysis of cognitive and literacy skills yielded several significant findings, revealing complex relationships between neurophysiological measures and reading abilities. These results contribute to our understanding of the neural underpinnings of reading proficiency and align with previous research in the field.

### Alpha Activity and Reading Proficiency

The low-reading group exhibited slightly higher peak alpha activity compared to the high-reading group, although these differences did not reach statistical significance. This observation is consistent with the complex relationships between alpha activity and cognitive performance reported in previous studies (Klimesch, 2012). The nuanced nature of this finding underscores the need for further investigation into the role of alpha oscillations in reading processes.

### Beta Power and Language Processing

A notable increase in beta power at frontal electrodes was observed in the high-reading group relative to the low-reading group. This finding corroborates the research of Weiss and Mueller (2012), which emphasized the critical role of beta oscillations in language processing and cognitive control. The enhanced beta power in skilled readers may reflect more efficient neural mechanisms for text comprehension and linguistic processing.

### Beta-Band Synchronization in Reading Networks

The high-reading group demonstrated enhanced beta-band synchronization in fronto-central-parietal networks compared to the low-reading group. This result aligns with the findings of Schurz et al. (2014), who highlighted the importance of these networks in efficient cognitive processing for skilled readers. The observed synchronization patterns may indicate more robust functional connectivity in brain regions crucial for reading comprehension and fluency.

### Alpha Band Associations and Developmental Perspectives

Complex relationships were observed in the alpha band, with higher age, grade level, better reading skills, and verbal fluency (measured by letters F, A, S) positively correlating with node centrality. Conversely, these factors negatively correlated with functional connectivity intensity in the alpha band. These intricate associations are consistent with developmental perspectives in neurocognitive research on reading (Franceschini et al., 2012). The findings suggest that as reading skills develop, there may be shifts in the functional organization of alpha band networks.

### Beta Band Associations and Developmental Trajectories

In younger participants, verbal fluency (A) showed positive associations with beta features in network power. Additionally, grade level positively correlated with peak alpha activity, while verbal fluency (A) negatively correlated with beta power. These results contribute to our understanding of the developmental trajectories of reading-related neural patterns, as discussed by Norton et al. (2015). The age-dependent associations highlight the dynamic nature of neural networks involved in reading acquisition and proficiency. In conclusion, these findings provide valuable insights into the neurophysiological correlates of reading ability, demonstrating the complex interplay between cognitive skills, age, and brain oscillatory patterns. Future research should further explore these relationships to elucidate the neural mechanisms underlying reading development and to inform educational interventions for improving literacy skills.

Therefore, those findings indicate notable neurophysiological differences between proficient and deficient readers, consistent with the growing body of research in reading neuroscience. The increased beta power in frontal regions among high readers aligns with existing literature suggesting that beta oscillations are crucial for language processing and cognitive control (Weiss & Mueller, 2012). This enhanced frontal activity may reflect more efficient cognitive processes in skilled readers, potentially facilitating better text comprehension and analysis. Enhanced synchronization within fronto-central-parietal networks among high-readers supports the notion that efficient reading relies on the coordinated activity of distributed brain regions (Schurz et al., 2014). The fronto-parietal network has been implicated in various cognitive functions, including working memory and attention, which are critical for successful reading (Sauseng et al., 2005). This finding suggests that proficient readers may have more integrated neural networks supporting their reading abilities.

The observed associations between alpha band measures and factors such as age and verbal fluency suggest that reading proficiency is linked to efficient neural organization within key brain regions. The complex relationship between these factors and functional connectivity intensity in the alpha band may indicate compensatory mechanisms or developmental changes affecting neural organization over time. This aligns with developmental perspectives in reading research, highlighting the dynamic nature of brain-behavior relationships in literacy acquisition (Franceschini et al., 2012).

The differential associations observed in the beta band, particularly in younger participants, underscore the importance of considering developmental factors when interpreting EEG data related to reading ability. These findings contribute to our understanding of the neural basis of reading proficiency and may have implications for the development of targeted interventions for individuals with reading difficulties.

## Conclusion

This exploratory study highlights significant differences in neural activity between good and poor readers as assessed through EEG. The findings suggest that proficient readers exhibit enhanced beta power and synchronization within critical networks associated with cognitive functions. These results align with and extend previous research in the field of reading neuroscience (Norton et al., 2015). Understanding these neurophysiological underpinnings can inform interventions aimed at improving reading skills among individuals facing literacy challenges. Future research should aim to replicate these findings with larger sample sizes and investigate the causal relationships between neural oscillations and reading ability through longitudinal studies and interventional approaches. The complex interplay between literacy, attention, and critical thinking skills emphasized in this study underscores the need for comprehensive approaches in education and cognitive development research. By bridging neuroscientific insights with educational practices, we can develop more effective strategies to support the development of essential cognitive skills across diverse populations.

